# Diotic and dichotic frequency discrimination thresholds in musicians and non-musicians: relationships between perception, musical ability and self-evaluated competence

**DOI:** 10.1101/810598

**Authors:** Gabriella Musacchia, Devin Inabinet, Jan De La Cruz, Justin Cha, Kevin Ng

**Author notes:** Corresponding Author: (GM). These authors contributed equally to this work (Gabriella Musacchia, Devin Inabinet, Jan De La Cruz). These authors also contributed equally to this work (Justin Cha, Kevin Ng).

## Abstract

Pitch perception provides important information for musical and vocal communication. Numerous studies have shown that musical training and expertise are associated with better pitch processing, however, it is unclear what types of pitch percepts are plastic with music training. The current study addresses this issue by measuring discrimination thresholds of Musicians (n=20) and Non-musicians (n=18) to diotic (same sound to both ears) and dichotic (different sounds to each ear) stimuli created from four types of acoustic computations:1) pure sinusoidal tones, PT; 2) four-harmonic complex tones, CT; 3) iterated rippled noise, IRN; and 4) interaurally correlated broadband noise, called “Huggins” or “dichotic” pitch sounds, DP. Frequency Difference Limens (DLF) in each condition were obtained via a 3-alternative-forced-choice adaptive task requiring selection of the interval with the highest pitch, yielding the smallest perceptible fundamental frequency (F0) distance (in Hz) between two sounds. Music skill was measured by an online test of musical Pitch, Melody and Timing (International Laboratory for Brain Music and Sound Research, https://www.brams.org/en/onlinetest/). Musicianship, length of music experience and self-evaluation of musical skill were assessed by questionnaire. Results showed musicians had smaller DLFs in all four conditions and that thresholds were related to subjective and objective musical ability. In addition, self-report of musical ability was shown to be a significant variable in group classification, suggesting that the neurobehavioral profile of musicians includes self-evaluation of musical competence.

## Introduction

Musical training is associated with better pitch encoding and perception (for review see (1)). In music, pitch is the quality that most strongly defines the perception of the melodic contour. Each note in a melody has a pitch that is related to the lowest frequency of an instrumental sound, called the fundamental frequency (F0). The perception of pitch, however, is not only evoked by musical sound, but can emerge from a wide variety of acoustic components (for review see (2)). A classic way to determine someone’s pitch perception ability is by measuring the smallest perceptible pitch change from a center frequency, called a *difference limen for frequency* (DLF) (3, 4). In general, normal-hearing listeners can perceive a change in as little as 2-3 Hz from a center frequency under optimal listening conditions (5). Musicians can detect even smaller pitch changes, sometimes so minute that the change is undetectable by otherwise normal-hearing non-musicians (2, 6, 7). Not surprisingly, increased acuity in musicians is not limited to musical sounds, but extends to perception and processing of speech (8), non-speech ((6) for review (9)) and non-native language sounds (10).

A prevalent hypothesis is that musical training improves auditory encoding mechanisms that give rise to pitch perception. However, the auditory system utilizes several mechanisms to encode pitch-related acoustics and it is unclear which ones are most improved with music training. One way the auditory system works is by representing the “temporal code” of a stimulus in which auditory neurons *phase-lock*, firing at a rate that matches the period, or frequency inverse, of a sound (for review see (11)). During temporal encoding, sounds trigger networks of neurons to compute and extract temporal patterns that can give rise to pitch perception (12, 13). Music practice and performance could activate and strengthen the temporal synchrony of these networks, thereby improving representation and higher-order computation acuity. Another mechanism to encode pitch, called “place code”, functions such that different frequencies activate discrete regions of the inner ear and subsequent nuclei, producing a tonotopic map of frequencies at each processing station (for review, see (14)). For example, the perception of pitch rises as the region of maximal activation on the basilar membrane moves closer to the base of the cochlea. Music training could generate more precise and definite tonotopic maps due to top-down modulation induced by the increased prevalence and relevance of sounds in the musician’s environment (15–17). Finally, a pitch perception can be generated by presenting different sound components to each ear, creating a *dichotic* (binaural) or combined estimation of the sound’s pitch (18, 19). Although the music-related mechanistic hypotheses are less prevalent for dichotic plasticity, it is reasonable to suggest that music training could increase the accuracy of communication between the left and right ears, particularly during azimuth (horizontal) localization tasks such as identification of instruments in an orchestra (20). To encode sound, the auditory system will use or integrate information gathered from each encoding strategy presented above, depending on what acoustic features are present in the stimulus.

The working hypothesis that motivated this study was that music training engenders plasticity in specific auditory encoding mechanisms. Particularly, we posited that sounds reliant on temporal encoding would be impacted the most because playing music requires considerable focus on sound timing. To test this, we measured DLFs in musicians and non-musicians using four different types of sounds with different pitch-related acoustics (Fig. 1). Creation of the sounds was inspired by work describing how different degrees of temporal, place and dichotic encoding mechanisms can be elicited in the auditory system (21). In order to assess musical skill, all participants took an online musical test for pitch, melody and timing measurements and filled out a questionnaire that probed duration of musical training and subjective self-reports of musical skill and listening habits.

**Figure 1.**
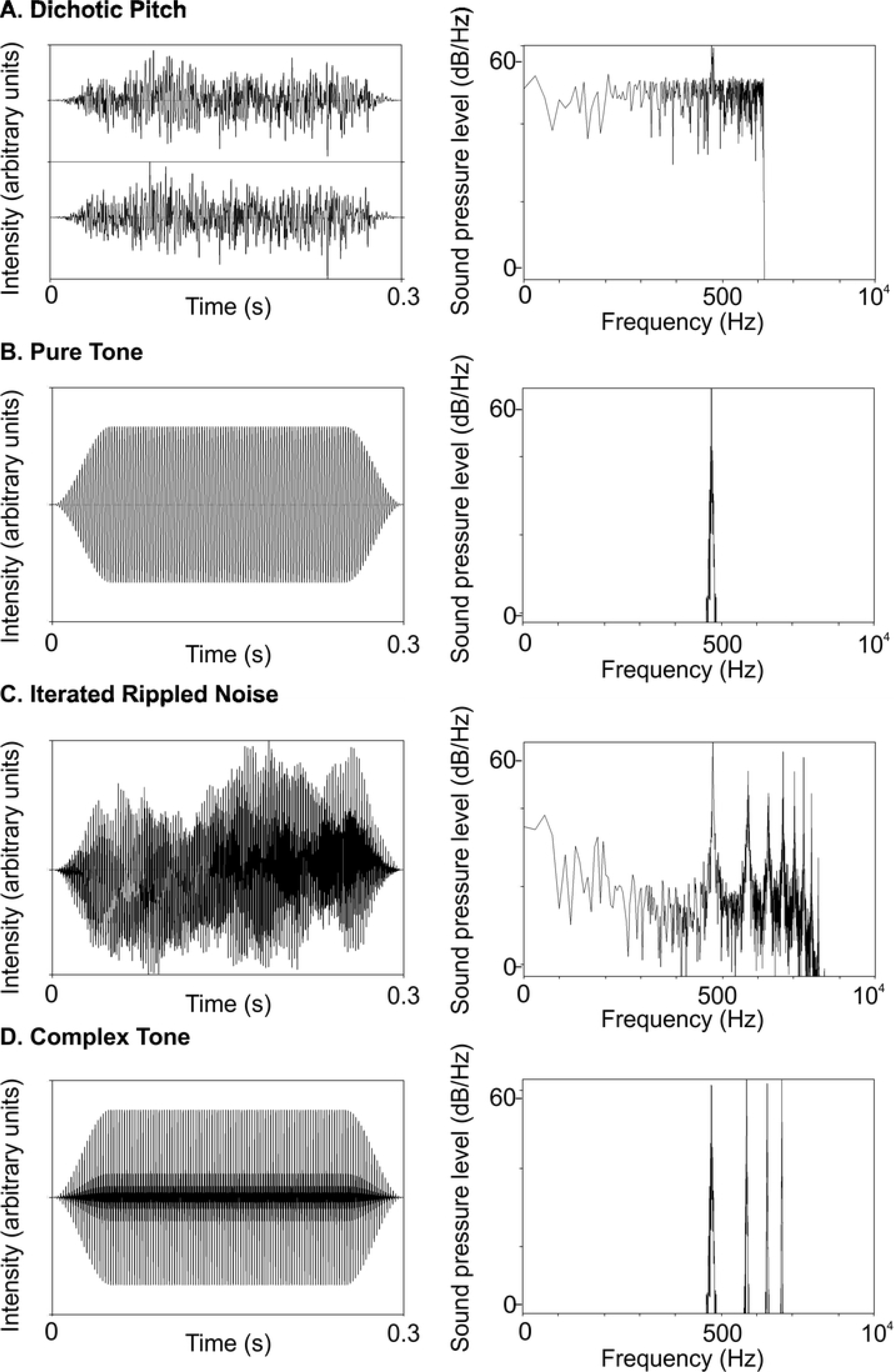
Study stimuli. Each row shows the 440 Hz stimulus waveform (right panel) and spectrum on a logarithmic frequency scale (left panel). A. Dichotic pitch with an interaural phase shift of 440 Hz, B. Pure Tone, C. Iterated Rippled Noise with a 64 iteration of delay and add at 1/440 s. D. Complex tone with three overtone harmonics

## Materials and Methods

### PARTICIPANTS

38 individuals with audiometric thresholds within normal limits (<25dB HL for 0.25, 0.5, 1, 2, 3, 4, 6 and 8 kHz) and no history of neurological disorders participated in the study. Previous research has shown that music-related brain plasticity is most effective when people begin playing music early, continue, and are currently practicing (22–24). Therefore, subject inclusion criteria in the Musician (MU) group included 1) self-identification as a musician via questionnaire and reported current involvement in musical activities, 2) self-report of music training initiation before high school (e.g. before grade 9, age 14-15) and 3) a total of at least 5 years in formal music education. 20 subjects fulfilled the criteria for MU group inclusion, with the remainder 18 subjects grouped into Non-musicians (NM). Group characteristics of age, music education, self-ratings and objective measures of musical skill (i.e. online aptitude test, for description see below) are presented in Table 1.

**Table 1.**
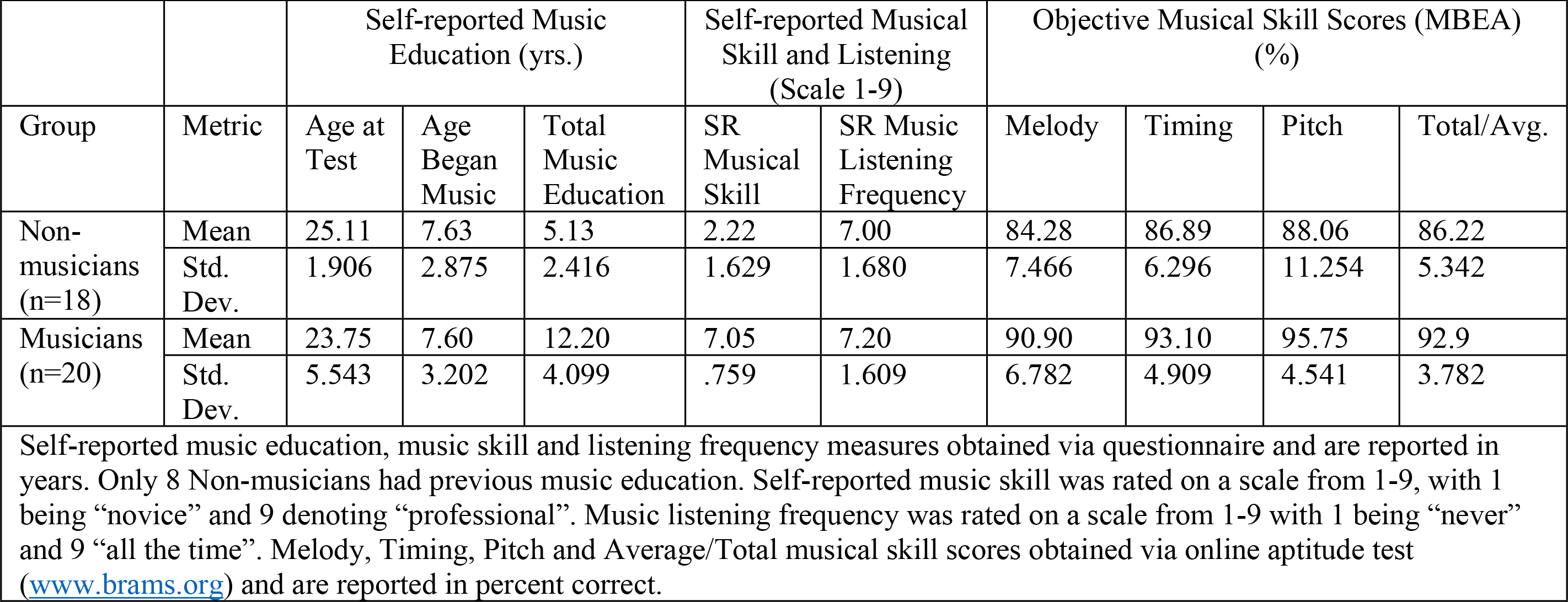
Group characteristics of music education, experience and skill in Non-musicians and Musicians

|  |  |  |  |  |  |  |  |  |  |  |
| --- | --- | --- | --- | --- | --- | --- | --- | --- | --- | --- |
| Group | Metric | Self-reported Music Education (yrs.) |  |  | Self-reported Musical Skill and Listening (Scale 1-9) |  | Objective Musical Skill Scores (MBEA) (%) |  |  |  |
|  |  | Age at Test | Age Began Music | Total Music Education | SR Musical Skill | SR Music Listening Frequency | Melody | Timing | Pitch | Total/Avg. |
| Non-musicians (n=18) | Mean | 25.11 | 7.63 | 5.13 | 2.22 | 7.00 | 84.28 | 86.89 | 88.06 | 86.22 |
|  | Std. Dev. | 1.906 | 2.875 | 2.416 | 1.629 | 1.680 | 7.466 | 6.296 | 11.254 | 5.342 |
| Musicians (n=20) | Mean | 23.75 | 7.60 | 12.20 | 7.05 | 7.20 | 90.90 | 93.10 | 95.75 | 92.9 |
|  | Std. Dev. | 5.543 | 3.202 | 4.099 | .759 | 1.609 | 6.782 | 4.909 | 4.541 | 3.782 |
| Self-reported music education, music skill and listening frequency measures obtained via questionnaire and are reported in years. Only 8 Non-musicians had previous music education. Self-reported music skill was rated on a scale from 1-9, with 1 being “novice” and 9 denoting “professional”. Music listening frequency was rated on a scale from 1-9 with 1 being “never” and 9 “all the time”. Melody, Timing, Pitch and Average/Total musical skill scores obtained via online aptitude test (www.brams.org) and are reported in percent correct. |  |  |  |  |  |  |  |  |  |  |

### STIMULI

Sounds were 300 ms in duration, with two 60-ms raised cosine ramps for onset and offset. Figure 1 shows time waveforms (left panels) and frequency spectra (right panels) for the 440 Hz (standard) stimuli used in the study. 440 Hz was chosen because it is a familiar musical note (A4) that elicits strong phase-locking.

In order to test binaural mechanisms, we created a dichotic pitch (DP) stimulus, often called “Huggins’ pitch,” which consists of dissimilar right and left inputs to make a dichotic estimation of a sound’s pitch (18, 19). DP stimuli were created with the Binaural Auditory Processing Toolbox for MATLAB8 using a transition width of 16%. DP sounds were made of white noise, diotic at all frequencies except for a narrow band at the F0 (440 Hz), over which the interaural phase transitioned progressively through 360°. Individuals were familiarized with DP perception through five online Demonstrations (https://web.stanford.edu/~bobd/cgi-bin/research/dpDemos/).

In contrast, a pure tone (PT), shown in panel B of Figure 1 is the product of a sinusoidal function. Sinusoids are thought to be encoded by place code mechanisms because they elicit narrow bands of maximal activation at specific places in the tonotopic map of the cochlea. At lower frequencies (<~2 kHz) elicit additional phase-locked temporal codes at the frequency’s period. Pure tones consisted of sinusoids at a fundamental frequency (F0) of 440 Hz, chosen.

We also tested an iterated noise (IRN) stimulus which evokes a pitch perception that is primarily reliant on temporal information (25–27). IRN stimuli, shown in Figure 1C, were created from Gaussian broadband noise filtered from 80-3000 Hz with 64 iterations of delay and add durations at the inverse of the F0 (440 Hz). The temporal regularity imposed on broadband noise gives rise to the perception of pitch despite low spectral content.

Finally, we used a complex tone with three harmonic overtones (CT), which most closely resembles the sound a musical instrument makes and relies on a combination of place and temporal codes. Complex tones (Fig. 1D) consisted of a four-harmonic complex (h1-h4) with equal amplitude and the same F0 (440 Hz).

### MUSICAL APTITUDE TEST

Individuals completed an online test through the International Laboratory for Brain, Music, and Sound Research (BRAMS, www.brams.org/en/onlinetest/) that allows for the assessment of the functioning of each musical component: 1) Melody, 2) Timing, and 3) Pitch ability. The online test battery is based on the Montreal Battery for Evaluation of Amusia (MBEA) and consists of musical phrases that vary along the melody, timing or pitch dimension (28). During the MBEA, listeners perform a two-alternative forced choice task to determine whether two presented musical phrases are the same or different. In the melody test, for example, the two choices may consist of an original melodic contour and a scale- or contour-violated alternate. The output of the online test is a percentage correct for each task, in addition to the average of all three categories. These percentages were recorded and utilized in this study. Group means and standard deviations are shown in Table 1.

### QUESTIONNAIRE

Participants’ musical history was collected through a questionnaire probing a range of information regarding subjective aptitude and measures of musicianship. We used the following details and scale-based ratings to correlate with objective performance on psychoacoustic measures: 1) Musician self-identification (e.g. “Are you a musicians?”), 2) Self-Report of Music Listening Frequency on a scale of 1-9 3) Self-Report of Musical Skill on a scale of 1-9 4) Age of Music Start and 5) Years of Consistent Practice. Group means and standard deviations are shown in Table 1.

### DATA ANALYSIS

Tests of normality were computed on all variables. Results of these tests showed that the pairs of MU and NM distributions were not significantly different from normal according to or Shapiro-Wilk tests (p<0.01), except for SR Musical Skill and BRAMS Pitch score (See Supplementary Table 1). Examination of the detrended SR Musical Skill scores showed that one NM rated themselves >1 Standard Deviation from normal and one MU rated themselves >−1 Standard Deviation from normal. Examination of the detrended BRAMS pitch scores showed that one individual from each group scored >−1 Standard Deviation from normal. Given that a skew in distribution was observed for two measures, we provide observed power for each test and only conducted tests that were robust to the assumption of normality (29, 30).

The group difference hypothesis was tested using a set of mixed repeated-measures ANOVAs (RMANOVA) for group (Musicians vs. Non-musicians, 2 between-subjects factors) and the following within-subjects factors: 1) sound type threshold (CT, PT, IRN, DP, 4 within-subjects factors), 2) sound type standard deviation (CT, PT, IRN, DP, 4 within-subjects factors), 3) self-reported measures of music skill and 4) listening frequency from the questionnaire (SR Musical Skill and SR Frequency of Music Listening, 2 within-subject factors). F-statistics, p-values and the observed statistical power, ranging from 0 to 1, where the fraction represents the chance of failing to detect an effect, are reported with each significant ANOVA result. Post-hoc t-tests were conducted when significant interaction effects with p-val<0.05 were observed.

To examine the question of a relationship between DLFs, musicality and self-assessment, Pearson’s r correlations were computed. Pearson’s r-values and p-values of the significance test are reported. In order to discover the degree to which our dependent variables discriminate between NM and MU, a discriminant function analysis with predictive classification of cases was conducted. The discriminant analysis included all four DLF thresholds, BRAMS total score and self-reported measures for a total of seven continuous, numeric variables and one categorical variable with two levels (NM, MU).

## Results

### Pitch discrimination thresholds

The 2X4 mixed RMANOVA on group (MU vs. NM, between-subjects factor) and threshold (CT, PT, IRN and DP, within-subjects factors) showed a within-subjects main effect of sound type; F(3,108)=38.137, p<0.001, Observed Power=1.0, an interaction effect F(3,108)=11.754, p<0.001, Observed Power=0.999, and a between-subjects main effect F(1,36)=29.205, p<0.001, Observed Power=1.0. Post-hoc t-tests were significant for all conditions (p<0.016).

Group means show that musicians had lower thresholds for each sound type category (Fig. 2, Supp. Table 1). Bar graphs in Figure 2 illustrates the group mean values for each sound type category, showing lower means for the musician group in all categories, relative to Non-musicians. Taken together, the data show that musicians can hear smaller pitch differences that Non-musicians in all four pitch-evoking sound type categories, with the greatest difference in the DP condition and the smallest difference in the CT condition.

**Figure 2.**
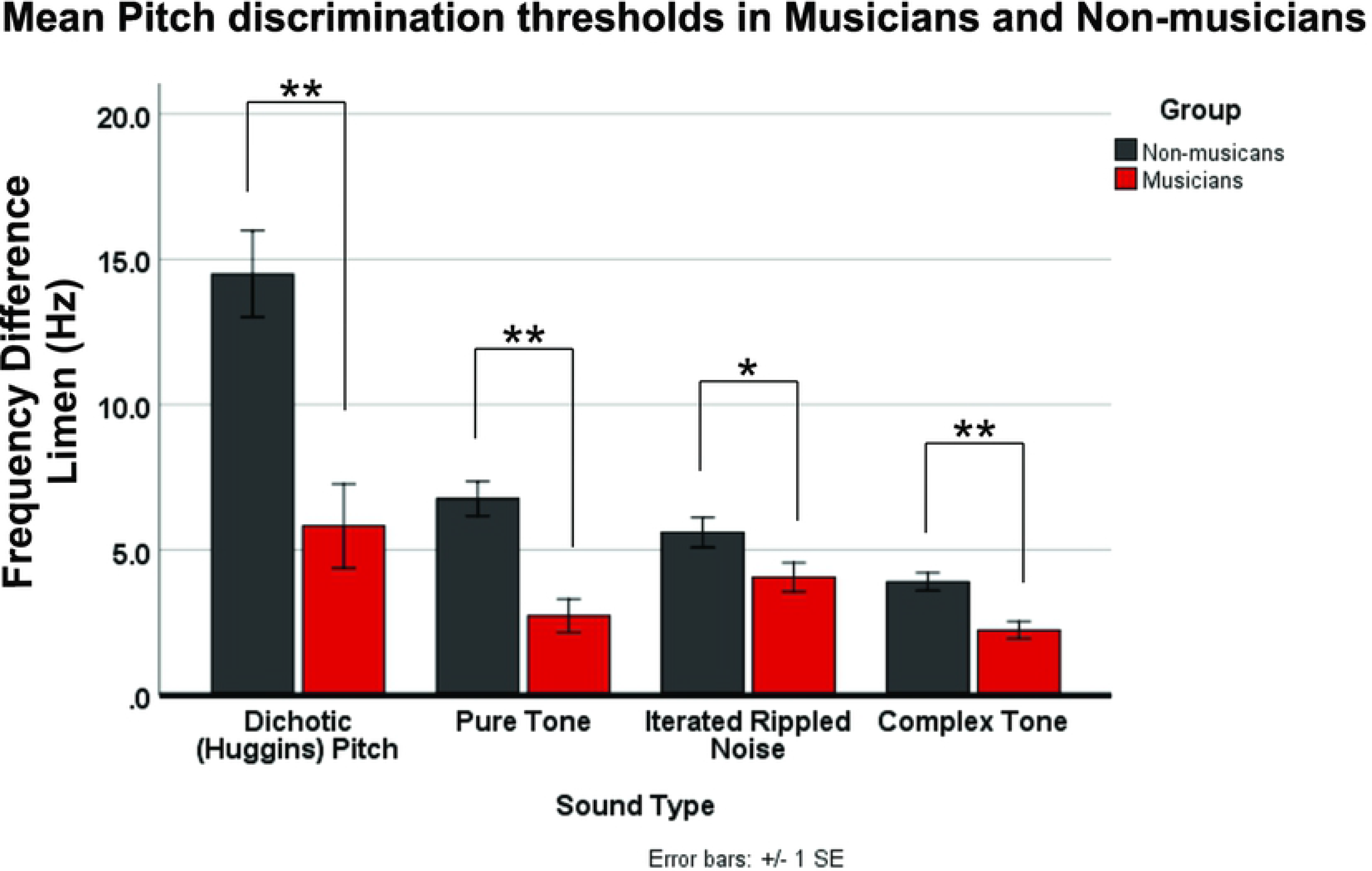
Bar graph shows mean DLF thresholds (+/− 1 SE). Musicians have smaller (better) pitch discrimination thresholds in all conditions, relative to non-musicians (*p<0.05; **p<0.01).

To examine group differences in threshold variance a 2X4 mixed RMANOVA on group (MU vs. NM, between-subjects factor) and threshold variance per sound type (DP, PT, IRN and CT, within-subject factors). For this analysis, standard deviation was computed from the four recorded threshold measurements per sound type. Results showed a within-subjects main effect of sound type; F(3,108)=18.265, p<0.001, Observed Power=1.0, an interaction effect F(3,108)=3.866, p=0.011, Observed Power=0.811, and a between-subjects main effect F(1,36)=11.702, p=0.002, Observed Power=0.914. Post-hoc t-tests were significant for DP and PT (p<0.007), but not IRN and CT (p>0.058). Examination of group means showed that threshold variance is smaller in MU than NM in the DP and PT condition (Supp. Table 2).

### Relationships between pitch discrimination thresholds, self-reports and musical aptitude measures

Pearson’s correlations show that better discrimination thresholds are associated with a higher self-report of musical skill and better scores on all tests of BRAMs musical aptitude. Correlations are reported in Supplementary Table 2. Figure 3 shows individual data for the representative correlations between DLFs, self-report and BRAMS total score. Figure 3 (left column) illustrates that lower (better) DLFs are associated with higher self-reports of musical skill. The spread of the data in Figure 3 also illustrates greater variance in self-report among NM compared to MU, reflecting a wider range of self-assessed musical experience in the NM group. Figure 2 (right column) shows the relationships between DLFs and BRAMS total score. Consistent negative correlations suggest that smaller DLFs are associated with higher musical aptitude.

**Figure 3.**
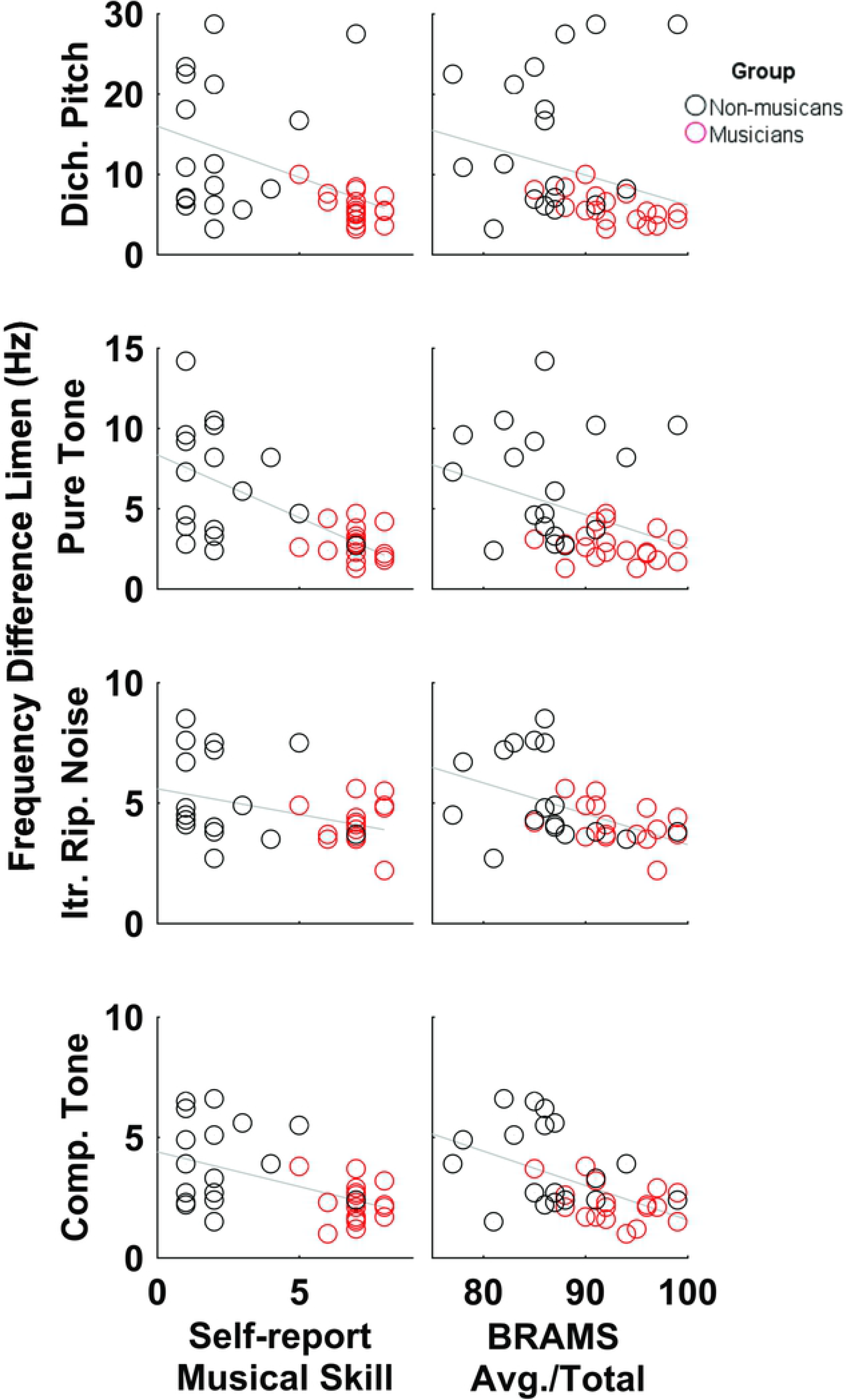
Scatterplots of individual data for Musicians (red) and Non-musicians (black) with regression lines. Left column shows relationships between pitch discrimination thresholds self-reported (subjective) musical skill (scaled between 1-9, with 1 being novice, 9 professional). Higher self-report is associated with smaller (better) thresholds. Right column shows relationships between pitch discrimination thresholds behavioral scores obtained from the BRAMS musical skills test (objective). Higher score is associated with smaller (better) thresholds.

### Discriminant analysis

A discriminant analysis was conducted to determine which of our variables contributed most to group separation and to test whether an individual’s group category could be correctly identified based on our continuous numeric experimental measures. Continuous variables were DP, PT, IRN and CT DLFs as well as SR Musical Skill and BRAMS Avg./Total score. Table 2 shows significant mean differences were observed for all variables (p<0.021) except for SR Music Listening Frequency (p=0.774).

**Table 2.**
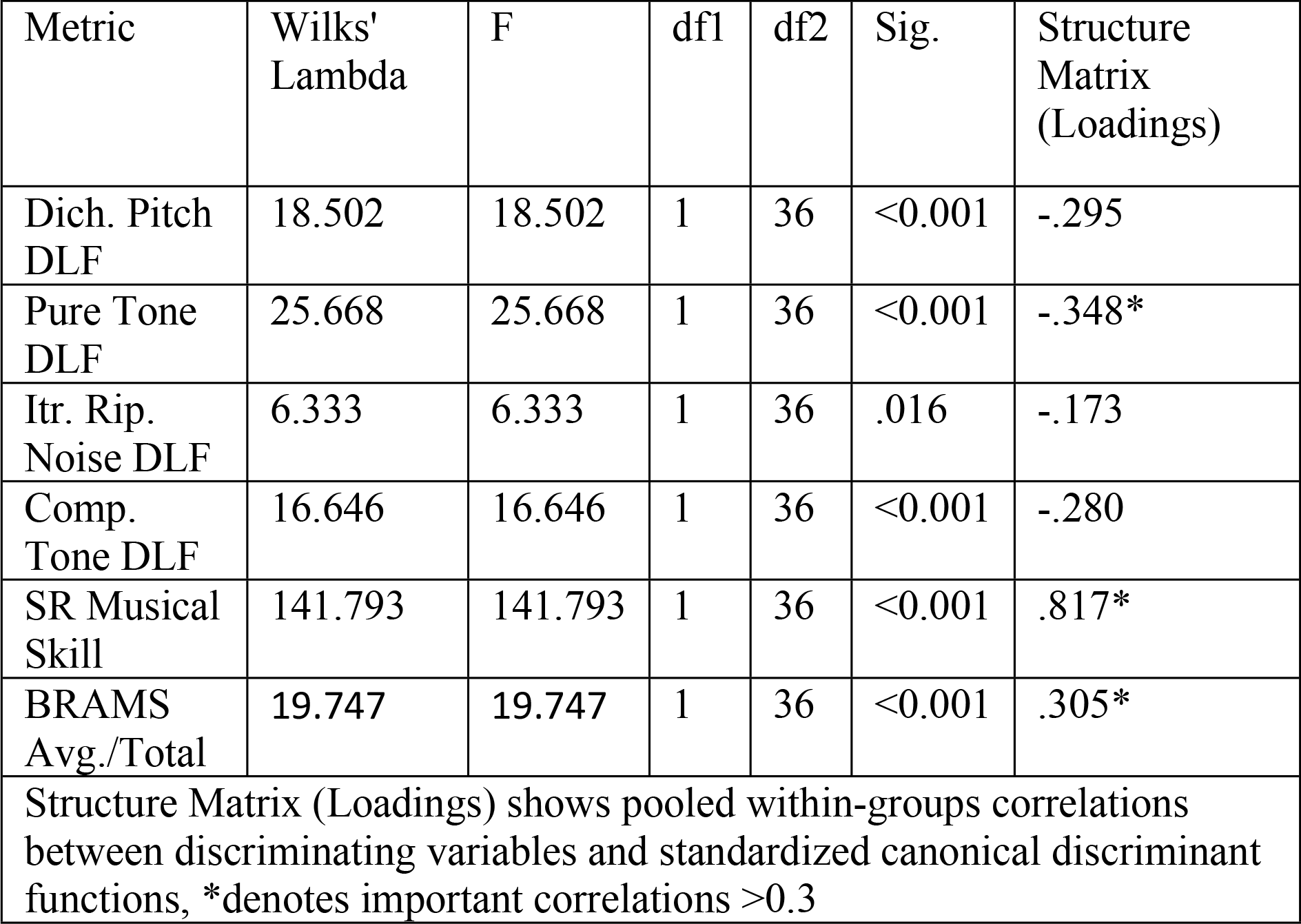
Discriminant analysis results including tests of equality of group means and variable loadings.

The canonical discriminant function showed a significant association between groups and variables; Wilks’ Lambda=0.145, Chi-square=60.800, p<0.001, accounting for 85.5% of the between-group variability. Examination of the discriminant loadings (Table 4) showed three significant predictors (i.e. >0.3), namely SR Musical Skill (.811) and PT DLF (−.343), and BRAMS Avg./Total score (.301). The weakest predictor was IRN DLF (−.169). Cross-validated classification showed that overall, 89.2% of the subjects were correctly classified into MU and NM groups. It should be noted that log determinants of this analysis showed large differences and Box’s M was significant, suggesting that the assumption of equality of covariance matrices was violated. However, this problem is somewhat allayed given that normality is not a critical assumption for discriminant analysis.

## Discussion

We have answered two main questions in this study: 1) Are musicians better at perceiving specific pitch-related acoustics? and 2) Are psychoacoustic thresholds related to objective and subjective measures of music ability? To answer these questions, we recorded psychoacoustic DLFs to four types of pitch-eliciting sounds in 18 Non-musicians and 20 Musicians and obtained objective and subjective measures of musicianship from all participants. Sound types for DLF measurement included Dichotic (Huggins’) pitch, Pure Tones, Iterated Rippled Noise and Complex Tones. Objective measures of musicianship were obtained from the output of the BRAMS online music aptitude test and subjective assessment of music ability and listening were obtained through the questionnaire.

To answer the first question, DLF data were subjected to a RMANOVA with four within-subject factors of sound type and two between-subject factors of group. Results showed group differences across all sound types, with the greatest differences for dichotic and pure tone stimuli. These data refute our initial hypothesis that pitch-related temporal encoding mechanisms would be most impacted by musicianship; instead suggesting that music-related plasticity is not restricted to types of pitches. The greatest difference between Musician and Non-musician discrimination thresholds in the dichotic condition suggests that higher-order mechanisms, such as those requiring a combination of sound across the ears, are greatly impacted by musical training.

Several hypotheses could reasonably explain our findings. One hypothesis is that mechanisms of music-related brain plasticity are not restricted to place or temporal code encoding mechanisms in peripheral or brainstem nuclei (11), but may also occur cortically (12), or at least beyond the superior olive where dichotic sounds first combine. Unfortunately, our current data do not permit further elucidation on the veracity of this postulate because we do not have encoding data to test brainstem and cortical plasticity specifically. An alternative hypothesis is that playing music sharpens one’s ability to extract pitch percepts in conditions where the pitch strength is less salient, such as the dichotic and iterated rippled noise conditions. If this hypothesis were true, we might expect that the largest differences between the two groups would be in the least salient conditions. Whereas the largest threshold difference is in the dichotic condition (less salient pitch), the second largest threshold difference is observed in the pure tone condition, which has the most salient pitch strength. Although our data do not directly address the issue of pitch strength, the fact that the largest differences are observed with both strong and weak pitch percepts diminishes this hypothesis’ likelihood. A third, big picture, hypothesis is that Musicians possess a greater aptitude to learn the task than Non-musicians. If this were true, we would expect Musicians to learn the task faster than Non-musicians. A post-hoc examination of the within-session change in threshold showed that Non-musicians did have more variability, measured by standard deviation (Supp. Table 4). However, mean magnitudes of the within-session change in threshold over the four runs, computed by subtracting the threshold obtained in the first run from the threshold obtained in the last run, did not appear to differ between groups (Supp. Table 4). To verify our observations, we performed two RMANOVAs for sound type and group on the standard deviation and within-session change data. Results showed that Musicians had lower standard deviation in thresholds to dichotic and pure tone stimuli, compared to Non-musicians, but only in the pure and complex tone conditions. No significant differences were observed for within- or between-subject comparisons of the within-session threshold change magnitude. Taken together, these data suggest that acclimatization or learning trajectories from task beginning to end is similar in Musicians and Non-musicians and that musicianship positively influences dichotic and pure tone pitch discrimination, in part by stabilizing threshold reliability.

In answering the second question, we showed evidence for a relationship between psychoacoustic pitch discrimination and measures of subjective and objective music ability. The correlation data show that discrimination thresholds across all four pitch types were negatively correlated with a higher subjective rating of musicianship, such that individuals who rated themselves with musical ability closer to “professional” on a subjective scale, could hear smaller pitch differences in all four sound conditions. Conversely, individuals who rated themselves with a lower musical ability (i.e. closer to “novice” on the same scale) had greater (poorer) DLFs. To the authors’ knowledge this is the first time that such a relationship has been reported. The results imply that a person’s self-assessment can be a good predictor of their psychoacoustic threshold. It should be noted however, that while the correlations between self-reported music ability and DLF are significant, more than half of the r-values portray a moderately strong relationship (i.e. <0.5); suggesting that other, untested, variables account for additional variance in the relationship. Therefore, while the connection between basic sensory ability and self-assessment of music ability is suggested here, it is only partially accounted for.

Regarding objective measures of music ability, it appears that the online music aptitude tests of Melody, Pitch, and the average of all music scores showed a consistent relationship with our psychoacoustic test results. In general, higher scores were related to smaller DLFs, suggesting that those who scored well on the BRAMS tests could discriminate sounds with smaller pitch differences. It is interesting to note that although Melody and Pitch scores correlated with psychoacoustic discrimination thresholds, Timing scores did not. Relatedly, Melody and Pitch were correlated to each other; r=0.419, p=0.010, but neither correlated with Timing; p>0.222. Timing scores did correlate, however, with SR Musical Skill; r=0.419, p=0.010, and BRAMS Avg./Total score; r=0.505, p=0.001, suggesting that rhythmic ability is related to musical aptitude and self-assessment of musical skill, but may be independent of pitch perception. Taken together, the correlation data show that the ability to discriminate small pitch differences can be reflected in global musical abilities and an individual’s evaluation of their own musical aptitude. This implies that sensory thresholds for pitch discrimination underlie, at least in part, one’s musical ability and self-appraisal of that ability. Furthermore, relationships between sensory threshold for pitch and more broad measures of musicianship are not restricted to a specific mechanism of pitch processing.

The Discriminant Analysis allowed us to detect the degree to which our variables discriminate between Musicians and Non-musicians. The variables that contributed most to the predictions of group membership were 1) Self-report of musical ability on a scale of 1-9, 2) Pure Tone DLFs and 3) BRAMS Avg./Total score. While the relationship between pure tone perception, musical aptitude and musicianship is well established, the contribution of a self-report variable is novel as far as the authors’ knowledge. Here, we show that self-evaluation of musical competence can be meaningfully applied to classify groups and is related to objective measures of music and perceptual ability. Self-evaluation of competence, or self-competence is defined as the sense of one’s capacity. (31) Previous data on this topic show that general self-competence is as associated with measures of cognitive ability such as IQ and academic achievement measured by GPA. (32) Our data support the argument that self-evaluation of competence is a meaningful measure of ability and outcomes (33) and extend into musicianship.

In addition to the finding of self-report as a meaningful measure, the discriminant analysis showed common characteristics of musicians include psychoacoustic, musical and self-evaluated abilities. This gives rise to the notion that all three areas may interact to define a person who is talented or skilled in music. It is interesting to note that the self-reported music listening scale did not distinguish between groups. This supports several lines of research showing that active music-making, rather than listening alone, is a catalyst for brain plasticity and internalized perceptual change (23, 34, 35).

In conclusion, this study sheds light on several aspects of musicianship. First, we show that the influence of musicianship is not limited to pitch judgements involving monotic/diotic mechanisms but also includes those that rely on dichotic integration. Second, our data show that basic perceptual thresholds are related to measures of both subjective and objective musical ability. And third, the data suggest that self-evaluation of musical ability is a meaningful part of musicianship such that high evaluation of competence are characteristic of musician group members. Taken together, the data update the neurobehavioral profile of musicians and extend creative ability measurements into new arenas.

## Acknowledgements

The authors gratefully acknowledge the class of University of the Pacific 2020 for participation in this study and Dr. Ganesh Swaminathan for his valuable comments on the manuscript. This work was supported by a 2019 Scholarly/Artistic Activities Grant (SAAG) from the University of the Pacific Faculty Research Committee.

## References

1. Munte TF, Altenmuller E, Jancke L. The musician’s brain as a model of neuroplasticity. NatRevNeurosci. 2002;3(6):473–8.

2. Micheyl C, Delhommeau K, Perrot X, Oxenham AJ. Influence of musical and psychoacoustical training on pitch discrimination. Hear Res. 2006;219(1–2):36–47.

3. Moore BC. Relation between the critical bandwidth and the frequency-difference limen. The Journal of the Acoustical Society of America. 1974;55(2):359.

4. Wier CC, Jesteadt W, Green DM. Frequency discrimination as a function of frequency and sensation level. The Journal of the Acoustical Society of America. 1977;61(1):178–84.

5. Moore JK, Linthicum FH, Jr. The human auditory system: a timeline of development. Int J Audiol. 2007;46(9):460–78.

6. Kishon-Rabin L, Amir O, Vexler Y, Zaltz Y. Pitch discrimination: are professional musicians better than non-musicians? J Basic Clin Physiol Pharmacol. 2001;12(2 Suppl):125–43.

7. Krishnan A, Gandour JT, Bidelman GM, Swaminathan J. Experience-dependent neural representation of dynamic pitch in the brainstem. Neuroreport.2009;20(4):408–13.

8. Musacchia G, Sams M, Skoe E, Kraus N. Musicians have enhanced subcortical auditory and audiovisual processing of speech and music. Proceedings of the National Academy of Sciences of the United States of America. 2007;104(40):15894–8.

9. Zatorre RJ. Neural Specialization for Tonal Processing. In: Zatorre RJ, Peretz I, editors. The biological foundations of music. Annals of the New York Academy of Sciences -- v. 930.: ANYAS; 2001. p. 193–210.

10. Wong PC, Skoe E, Russo NM, Dees T, Kraus N. Musical experience shapes human brainstem encoding of linguistic pitch patterns. Nat Neurosci. 2007;10(4):420–2.

11. Pickles JO. The auditory nerve. In: Pickles JO, editor. An Introduction to the Physiology of Hearing. Fourth ed. Lieden, The Netherlands: Brill; 2013. p. 73–83.

12. Cariani PA. Temporal codes and computations for sensory representation and scene analysis. IEEE TransNeural Netw. 2004;15(5):1100–11.

13. Tramo MJ, Cariani PA, Koh CK, Makris N, Braida LD. Neurophysiology and neuroanatomy of pitch perception: auditory cortex. Ann N Y Acad Sci. 2005;1060:148–74.

14. Pickles JO. Physiological correlates of auditory psychophysics. In: Pickles JO, editor. An Introduction to the Physiology of Hearing. Fourth ed. Lieden, The Netherlands: Brill; 2013. p. 278–81.

15. Zhang Y, Suga N. Modulation of responses and frequency tuning of thalamic and collicular neurons by cortical activation in mustached bats. JNeurophysiol. 2000;84(1):325–33.

16. Bergan JF, Ro P, Ro D, Knudsen EI. Hunting increases adaptive auditory map plasticity in adult barn owls. JNeurosci. 2005;25(42):9816–20.

17. Coffey EBJ, Musacchia G, Zatorre RJ. Cortical Correlates of the Auditory Frequency-Following and Onset Responses: EEG and fMRI Evidence. The Journal of neuroscience : the official journal of the Society for Neuroscience.2017;37(4):830–8.

18. Cramer EMH, W.H. Creation of pitch through binaural interaction. The Journal of the Acoustical Society of America. 1958(30):412–7.

19. Yost WA. Thresholds for segregating a narrow-band from a broadband noise based on interaural phase and level differences. The Journal of the Acoustical Society of America. 1991;89(2):838–44.

20. Hairston WD, Hodges DA, Burdette JH, Wallace MT. Auditory enhancement of visual temporal order judgment. Neuroreport.2006;17(8):791–5.

21. Walker KM, Schnupp JW, Hart-Schnupp SM, King AJ, Bizley JK. Pitch discrimination by ferrets for simple and complex sounds. The Journal of the Acoustical Society of America. 2009;126(3):1321–35.

22. Musacchia G, Strait D, Kraus N. Relationships between behavior, brainstem and cortical encoding of seen and heard speech in musicians and non-musicians. Hear Res. 2008;241(1–2):34–42.

23. Kraus N, Strait DL. Emergence of biological markers of musicianship with school-based music instruction. Ann N Y Acad Sci. 2015;1337:163–9.

24. Skoe E, Kraus N. A little goes a long way: how the adult brain is shaped by musical training in childhood. JNeurosci. 2012;32(34):11507–10.

25. Yost WA. Pitch of iterated rippled noise. The Journal of the Acoustical Society of America. 1996;100(1):511–8.

26. Yost WA, Patterson R, Sheft S. A time domain description for the pitch strength of iterated rippled noise. The Journal of the Acoustical Society of America. 1996;99(2):1066–78.

27. Bidelman GM. Sensitivity of the cortical pitch onset response to height, time-variance, and directionality of dynamic pitch. Neurosci Lett. 2015;603:89–93.

28. Peretz I, Champod AS, Hyde K. Varieties of musical disorders. The Montreal Battery of Evaluation of Amusia. Ann N Y Acad Sci. 2003;999:58–75.

29. Tiku ML. Power function of the F-test under non-normal situations. Journal of theAmerican Statistical Association. 1971(66):913–5.

30. Tan WY. Sampling distributions and robustness of t, f and variance-ratio in two samples and ANOVA Models with respect to departure from normality. Communications in Statistics - Theory and Methods. 1982(11):2485–511.

31. Tafarodi RW, Swann WB, Jr. Self-liking and self-competence as dimensions of global self-esteem: initial validation of a measure. J Person Assess. 1995;65(2):322–42.

32. Mar RA, DeYoung CG, Higgins DM, Peterson JB. Self-liking and self-competence separate self-evaluation from self-deception: associations with personality, ability, and achievement. J Pers. 2006;74(4):1047–78.

33. Baumeister RF, Campbell JD, Krueger JI, Vohs KD. Does High Self-Esteem Cause Better Performance, Interpersonal Success, Happiness, or Healthier Lifestyles? Psychol Sci Public Interest.2003;4(1):1–44.

34. Musacchia G, Ortiz-Mantilla S, Choudhury N, Realpe-Bonilla T, Roesler C, Benasich AA. Active auditory experience in infancy promotes brain plasticity in Theta and Gamma oscillations. Dev Cogn Neurosci. 2017;26:9–19.

35. Gerry D, Unrau A, Trainor LJ. Active music classes in infancy enhance musical, communicative and social development. DevSci. 2012;15(3):398–407.

